# Homology-aware cross-validation strategies for generalization assessment in RNA secondary structure prediction

**DOI:** 10.64898/2026.06.28.735057

**Authors:** Leandro Bugnon, Guillermo Kulemeyer, Matias Gerard, Leandro Di Persia, Georgina Stegmayer, Diego H. Milone

## Abstract

RNA secondary structure prediction is a fundamental challenge in bioinformatics, essential for understanding the functional roles of non-coding RNAs. Recently, deep learning models have transformed the field with impressive results, leading to critical discussions regarding the validity of current crossvalidation strategies. On the one hand, traditional random partitioning yields overoptimistic results due to data leakage from uncontrolled homology. On the other hand, removing every training sequence that even slightly resembles the testing set penalizes learning-based methods, requiring generalization to completely out-of-distribution sequences. While removing sequences and retraining a machine-learned model is straightforward, the experimental data used for development and parameter tuning of classical thermodynamic methods cannot be removed, leaving these methods to benefit from knowledge leakage. Here we critically analyze the cross-validation strategies currently used for RNA secondary structure prediction: random splitting, clustering-based splitting, and leaving one RNA family out for testing. We demonstrate that this last strategy can leave structural patterns present only in the test partition, making the learning task impossible. Overall, we find that the apparent superiority of either type of method is an artifact of the splitting strategy. We propose here two homology-aware strategies: human-learned folds, which exclude from testing the structures already captured by decades of human learning, and similarity folds, which measure performance across the full spectrum of test-to-train similarity. Both enable fair comparisons, applying the same rigorous assessment to classical and learning-based methods.

Data and source code are available at https://github.com/sinc-lab/xvalRNAfolding

## Introduction

Ribonucleic acid (RNA) plays a crucial role in many fundamental biological processes, such as gene expression, cell signaling, and post-transcriptional regulation^1^. There is currently a wide variety of publicly available RNA sequences, and they keep growing at an ever-increasing rate^2^. Although the RNA structure is key to inferring their functions and understanding the underlying mechanisms, most of those structures remain unknown because their experimental determination is costly and time consuming^3, 4^. Moreover, RNA structures represent less than 1% of the total cataloged protein structures^5^, and their energy landscape is dominated by the secondary structure^6, 7^. To address these limitations, computational methods for secondary structure prediction have been proposed in the last decades, but the challenge is still one of the major and long-lived problems in computational biology^8–11^. Currently, state-of-the-art methods can be classified into two main categories: classical methods based on optimization with thermodynamics models; and deep learning (DL) methods.

In the last few years, while models like AlphaFold^12^ have provided high accuracy, boosting research in protein biology, the prediction of RNA structures with DL lags far behind^7^. The major challenge is the tendency of large models to memorize training data rather than learning fundamental underlying patterns. Furthermore, these models are often susceptible to shortcut learning^13^, where they achieve high performance by identifying unintended statistical associations or local motifs that do not reflect the complex biological rules of RNA folding. The increasing number of recently published DL models for RNA secondary structure prediction reporting impressive results has raised warnings about overstated performance^14, 15^. Thus, proposals for more rigorous cross-validation methodologies and better control of homology among data partitions have arisen in the community^16–18^.

A long-established procedure is partitioning the available data into different sets for different purposes: the train set for fitting the model parameters, the validation set for hyperparameter tuning and model selection, and the test set for estimating its future performance on unseen data^19^. Thus, train and validation sets are used during model development, while the test set is only used for the final evaluation of the model. The data partition into these subsets should ensure a truthful evaluation of the generalization capacity of the model, without being negatively affected by its memorization capacity^20^. For RNA secondary structure prediction, there are three splitting strategies widely used by the community. The most common one is the simple *k*-fold split, which randomly separates data samples into *k* train/test partitions, based on the assumption that data is equally distributed^21^. However, in the case of RNA folding, there are different RNA families with members that are very similar in sequence and structure. Thus, in a random partitioning the family members within the training and testing sets will intersect and reported performance will not adequately estimate the generalization capability to predict novel RNA structures, inflating results. A second strategy involves the splitting of datasets based on the sequence similarity between train and test sets^22–24^, for example by using a sequence-identity cutoff with CD-HIT^25^ or similar tools^26^. This strategy, hereafter referred to as cluster-fold, consists of grouping similar sequences into clusters, which are then assigned either to the train or test partition, thereby avoiding similar samples appearing in both partitions. However, since it relies on a single predefined cutoff between train and test, the resulting performance is highly dependent on this threshold. The third widely used method is cross-family validation^16^, hereafter referred to as fam-fold. In this approach, partitions are created such that there is no intersection between families: a complete RNA family is left out for testing, and the remaining families are used for training. This eliminates most of the homology to the training set, which presumably allows the estimation of future performance on never seen families. However, fam-fold generates partitions with a highly variable difficulty that cannot be controlled, and a large imbalance in the test partitions due to the inherent sizes of known RNA families. Moreover, as will be shown in this study, fam-fold can generate partitions in which the patterns to be learned do not exist in the training data.

In this work, we propose two additional methodologies for performance comparison that enable a rigorous and fair evaluation of both DL and classical models. In DL models, humans build a software with rules to be trained automatically from domain examples; therefore excluding those examples for testing is straightforward. This model is learned from data by a computational algorithm, that is, it is a machine-learned (ML) model. Conversely, in classical models, researchers themselves learn from data samples and then build a specific software tool based on the rules they learned^27^. This model is developed and fitted over decades of research, constituting a human-learned (HL) model. Excluding prior knowledge for testing is much more challenging in this context, as it would require removing from the test set all sequences that contributed to the expert knowledge used to produce the rules encoded in the model. We address this issue by proposing an analogous methodology to exclude the well-known sequences from the test set. Additionally, we show that an improved variant of cluster-fold partitioning can be used to explore the full spectrum of sequence similarities. This strategy provides a more comprehensive evaluation of generalization and fair comparison of methods across a wide range of similarity conditions between train and test partitions.

When reviewing RNA folding methods, it is common in practice that every new prediction tool uses its own dataset and its own validation schema^28^. Furthermore, most of the time, the ML models are not re-trained under the same conditions as the other competitors^11^, resulting in unfair comparisons (some exceptions are^29, 30^). Differently from other works, we provide here a practical setup to compare existing cross-validation methodologies and analyze them based on experimental validations with state-of-the-art folding methods. Although we acknowledge that the field is shifting towards 3D structure prediction, this study is specifically intended for RNA secondary structure prediction methods. Nevertheless, the key findings regarding the assessment of generalization capacity, and the discussion on data leakage versus knowledge leakage, could be experimentally extended to support 3D structure prediction methods in future work.

## Cross-validation strategies in RNA secondary structure prediction

### Random folds

The random *k*-folds strategy is based on the assumption that data is independent and identically distributed^21^. Fig. 1.a (top) depicts this methodology schematically. The data are randomly separated into *k* groups and then, at each fold, one group is selected as the test partition and the rest form the training partition. However, existing available biological databases for benchmarking are full of duplicates and highly similar sequences (homologs), due to the evolutionary relationships that exist in their generation, and the redundancy in experimental collection^31^. Thus, similar data points are spread among the train and test sets when created by random splitting, inducing data leakage^20^. Therefore, the main problem with this approach is that DL models can solve the predictions on the test sets by simple memorization or shortcut learning, leading to an overestimation of their predictive performance. In fact, in the last few years, a number of models based on DL reported impressive results for RNA secondary structure prediction^24, 32–35^. However, many of these works assessed performance using *k*-folds cross-validation. Since the structure of RNA is highly conserved inside a family, performance derived from this methodology does not demonstrate generalization to novel RNAs^36^.

**Fig. 1.**
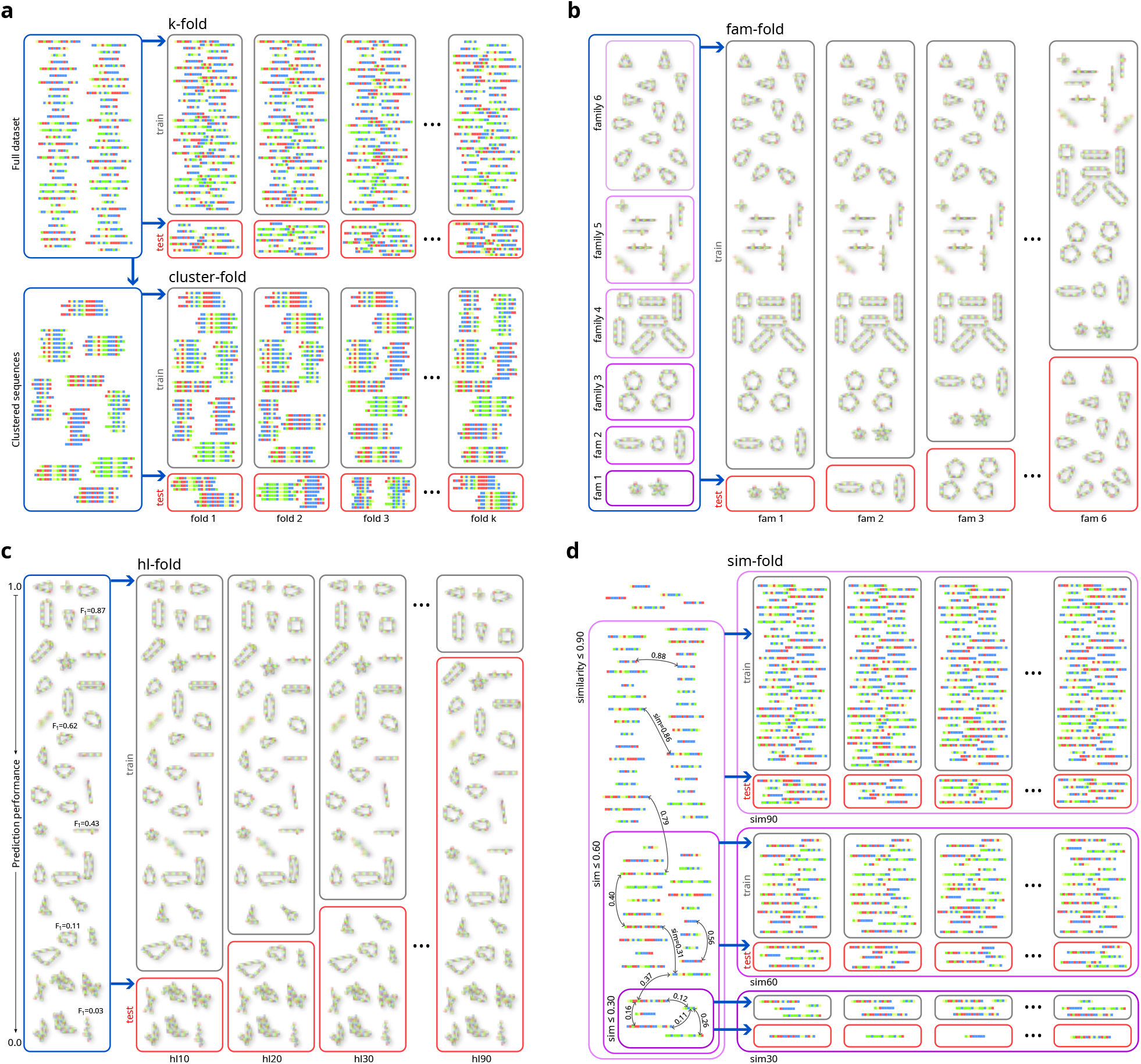
Cross-validation methodologies for RNA secondary structure prediction. **a** Random **k-fold** (top): the complete dataset of RNA sequences is randomly divided by *k* groups. In each fold, a group is used for testing (red) and the rest for training partitions (gray). In **cluster-fold** (bottom), the complete dataset is split into clusters of similar sequences. Then, for each fold a subset of these clusters is assigned to the training set, while the remaining clusters constitute the testing partition. **b fam-fold**: the illustration has 6 structural families (triangles, lines, ovals, etc.). At each fold, one complete family is left out and used only for testing, while all the other families are used for training; **c hl-fold**: each fold has in the training set all the sequences for which classical methods obtained an *F*_1_ *> τ* threshold, and the rest of the sequences are used for testing. Several values of *τ* are defined to build the folds. **d sim-fold**: several groups of increasing *σ* sequence similarity are built; then, inside each group of controlled similarity, many random train/test folds can be sampled.

### Clustering folds

This strategy addresses the data splitting into clusters of similar sequences (Fig. 1.a, bottom). Then, clusters are distributed among train/test partitions in each fold, preventing similar sequences from being in different partitions. This strategy removes data leakage up to a threshold of similarity among sequences^20^. It is worth noting that the most frequent practice with this strategy is to build only one fold, with the corresponding training and testing partitions.

For measuring similarity, pairwise alignment methods are used, particularly due to their easy interpretation. However, these methods have a high computational complexity, especially as the length of the sequences grows. In order to address large datasets, most of the clustering strategies for biological sequences attempt to reduce the number of alignments by using the well-established tools like Cluster Database at High Identity with Tolerance (CD-HIT)^25^. Partitions are generated subject to a single maximum level of similarity between train and test. A well-defined criterion to set up the allowed level of similarity is required. An example of cluster-fold can be found in^20^, which proposes SpanSeq, an alignment-free approach to data partitioning for DL applications, based on three steps: similarity calculation among all the sequences of the dataset, clustering of similar sequences into groups, and distributing them into *k* folds, minimizing the difference of the amount of samples between train and test partitions. In this work, a single threshold of similarity was set based on RNA bibliography. However, as the authors stated, this value might not be sufficient, so data-leakage might have occurred. Similarly, RNA3DB^37^ presents a dataset designed for benchmarking DL models for 3D RNA structure prediction, which provides coverage of all RNA chains found in the Protein Data Bank (PDB). Partitioning proceeds in stages: a sequence-based clustering, where all RNA chains with high sequence similarity are grouped and the longest RNA chain is chosen as cluster representative; and a structure-based clustering, where an RNA homology search against Rfam families is used to build a graph whose maximally connected components are guaranteed to share no structural homology. Both stages operate at a single, fixed operating point (99% identity and an Infernal E-value cutoff of 1.0, deliberately set to a not strict value so that no potential homology is missed), and the released dataset is then a single train/test split in an approximate 70/30 ratio. Although alternative splits can be obtained by changing parameters, no multi-threshold partitioning is reported. In CASP-like time-based evaluations of RNA^38^, the test sequences are collected over time following the submission deadline, after which the methods cannot be updated. Up to that date, methods may be trained with any dataset publicly available. However, in this case the similarity between test and train is not controlled and mixes several similarity levels. Overall, in practice multi-threshold similarity-based evaluation is rarely performed in RNA secondary structure prediction literature, as recent reviews indicate^14, 15^. Thus, the main problem with the practical use of cluster-fold approaches is that the criteria for defining the cut-off are unclear in most of the tasks, and the performance obtained is highly dependent on the selected threshold, with no guarantee of avoiding data-leakage in the commonly used greedy algorithms.

### Family folds

Another method proposed for cross-validation in RNA secondary structure prediction is the named interfamily or fam-fold, firstly proposed in^16^. This methodology proposes that one family is left out for testing per cross-validation fold. That is, 8 families are used for training, leaving 1 family for testing in each fold. This is done for every family in the dataset. Fig. 1.b illustrates this strategy with 6 schematic RNA families. The motivation behind this is to measure model performance on novel RNAs that do not belong to a known family. This promises to eliminate most of the homology to the training set, providing a hard measure of generalization and, thus, allowing estimation of future performance on novel RNAs that do not belong to any well-known family.

The problem with this strategy is twofold. On the one hand, classical methods will always obtain acceptable performance in the fam-fold setup because they use constraints and thermodynamic parameters that have been experimentally determined from real hairpins and other important structures frequently found in all RNA families^39–43^. This can be considered a form of data leakage —or more specifically, knowledge leakage— as information from the test set could be implicitly utilized to design the rules of the model or fine-tune its underlying parameters. On the other hand, fam-fold eliminates from the training set key patterns that are present only in the test family, which is unfair for data-driven methods. In^44^ it was already demonstrated how performance significantly declines when DL algorithms are trained on a specific set of families or sequence lengths and then tested on different ones. Fam-fold strategy tries to avoid overestimating performances by simple memorization. However, if there are certain structural patterns that are exclusive to a family, and assuming that the DL model has the capacity to learn them, removing that family from the training set violates the basic hypothesis of ML —namely, that a model cannot learn from patterns that do not exist within the training data. These particular patterns of a RNA family may even be the result of different protocols and experimental conditions under which their reference structures were obtained, since they come from independent studies. Thus, there might not be a single thermodynamic model that covers all cases based solely on sequence information. This also explains why thermodynamic methods are far from providing a definitive prediction, with very low performance for some families like tmRNA.

## Controlling homology levels

### Human learned folds

One of the ways in which human beings understand and learn from nature is by developing conceptual models that can explain the reality observed through data. Then, the use of these models on new data can be automated through the implementation of a software tool. From this perspective, computational models can be classified into “machine-learned”, where humans build a software with rules to be trained automatically from domain examples; and “human-learned”, where humans learn themselves from data samples and then build a software with those rules provided by domain experts. On the one hand, in rigorous homology-aware validations of RNA secondary structure prediction, ML models still provide just similar or slightly lower performance than the HL ones^16, 30^. Moreover, they have other disadvantages: those are hard to interpret by humans, hard to realize how features interact, and the model does not provide an easy understanding of the nature of the data (and the phenomena) under analysis^45^. On the other hand, a HL model is developed standing on previous theories (for example, thermodynamics), and it is built on case by case studies. This “training” is done over many years, even decades, because the human learning process is slow. These models have the main advantage of providing interpretable explanations of the processes that take place, i.e. a mechanistic model based on previous knowledge, which in turn can give rise to a novel theory^46^. HL models enable relatively simple cause-effect analysis that does not rely on large amounts of data, because actually it is the human being who has seen many examples and has come up with a reduced set of hypotheses to test the law behind them. The disadvantage is low predictive capacity when there is an inherent complexity in the data or the processes involved in its generation. Moreover, these models are based only on very well-known and well-studied facts. Those are mainly fitted to the known and not prepared for the discovery of the new.

From a learning perspective, while hacking a machine-learned model is relatively simple, by removing homologs from the train and test partitions, it is really a challenge to do such a similar splitting for an HL model. It can be stated that the accumulated human knowledge over the last 40 years, used to create a specific tool, is the training set for the HL models. In the case of RNA folding, RNAfold^47^ is one of the most widely used HL methods, based on classic thermodynamics and optimization algorithms^48^. This model has been adjusted over decades based on experimental observations, biochemical and thermodynamic theories, errors observed from the model itself, case-by-case studies of sequences over several years, adjustments of parameters, etc. Thus, the best predicted structures of this HL model can be considered as representatives of the training partition, that is, the sequences for which human knowledge has been able to learn and build a proper model. Taking all this into account, we propose to define several hl*τ* -folds by considering all the sequences for which a consensus of classical methods (RNAfold, RNAstructure, LinearFold and LinearPartition) provides a performance higher than a given threshold *τ*, and the rest of the sequences are left for testing. Fig. 1.c illustrates this strategy. For example, the hl70-fold has, in the train partition, all the sequences for which the consensus mean of *F*_1_ is above the threshold of 0.70, and the rest of the sequences are in the test partition. In this way, all the structures that have been well learned and incorporated into the HL methods are not considered for testing, because they were evidently part of the training set used for human learning throughout the last decades. Thus, by using different values of *τ*, a spectrum of difficulties can be obtained to evaluate the prediction methods.

### Similarity folds

As stated for cluster-fold, data splitting into groups of similar sequences avoids data leakage, but just up to a certain (single) point of similarity. This strategy requires choosing a well-defined criterion to set up the allowed level of similarity between train and test partitions. Instead, a better approach should be taking into account the full spectrum of possible similarity sub-sets existing within the dataset under analysis, in order to provide a real evaluation of generalization to a wide range of similarity levels between testing and training partitions.

Therefore, we propose here the sim-fold splits to explore the full-range of similarities within any dataset as a generalization and formalization of a multi-threshold similarity-based evaluation. It is a framework for comprehensive model evaluation that plots model performance as a function of increasing test-to-train similarity levels. This strategy is schematically shown in Fig. 1, panel d). Available data are grouped into subsets, where the similarity among all sequences in the subset is below a given maximum. Multiple subsets sim*σ* are created by applying similarity thresholds *σ* from 10 to 90%. For example, the sim30 split will be composed of all RNA sequences that have a maximum of 30% similarity among them. Then, inside each sim*σ* split, several folds can be made randomly with the corresponding traintest partitions. Since maximum similarity is ensured among all sequences of a given sim*σ* split, the maximum similarity between test and train partitions is guaranteed for any random sampling of that split. Overall, across all similarity thresholds the sim-fold strategy allows a comprehensive evaluation that plots model performance as a function of increasing similarity between testing and training data.

## Multi-threshold splitting strategies are key for properly estimating generalization capability

Figure 2 shows the performance of the RNA secondary structure prediction methods according to different cross-validation strategies in the Archive II dataset. It shows, for each strategy, the distributions of the test-to-train structural distances of each fold, computed using RNAdistance from the ViennaRNA package^47^, the *F*_1_ scores for each prediction method, and the relative size of the test partitions of each fold. The DL prediction methods include REDfold, UFold, sincFold, MXFold2 (hybrid), and the RNA foundation model RiNALMo^49^, indicated as the best one in a recent review of RNA large language models for secondary structure prediction^30^. All DL and hybrid models were retrained from scratch for every fold in each cross-validation strategy. The classical prediction methods included are RNAfold, RNAstructure, LinearFold, LinearPartition, ProbKnot and IPknot (details in Methods).

**Fig. 2.**
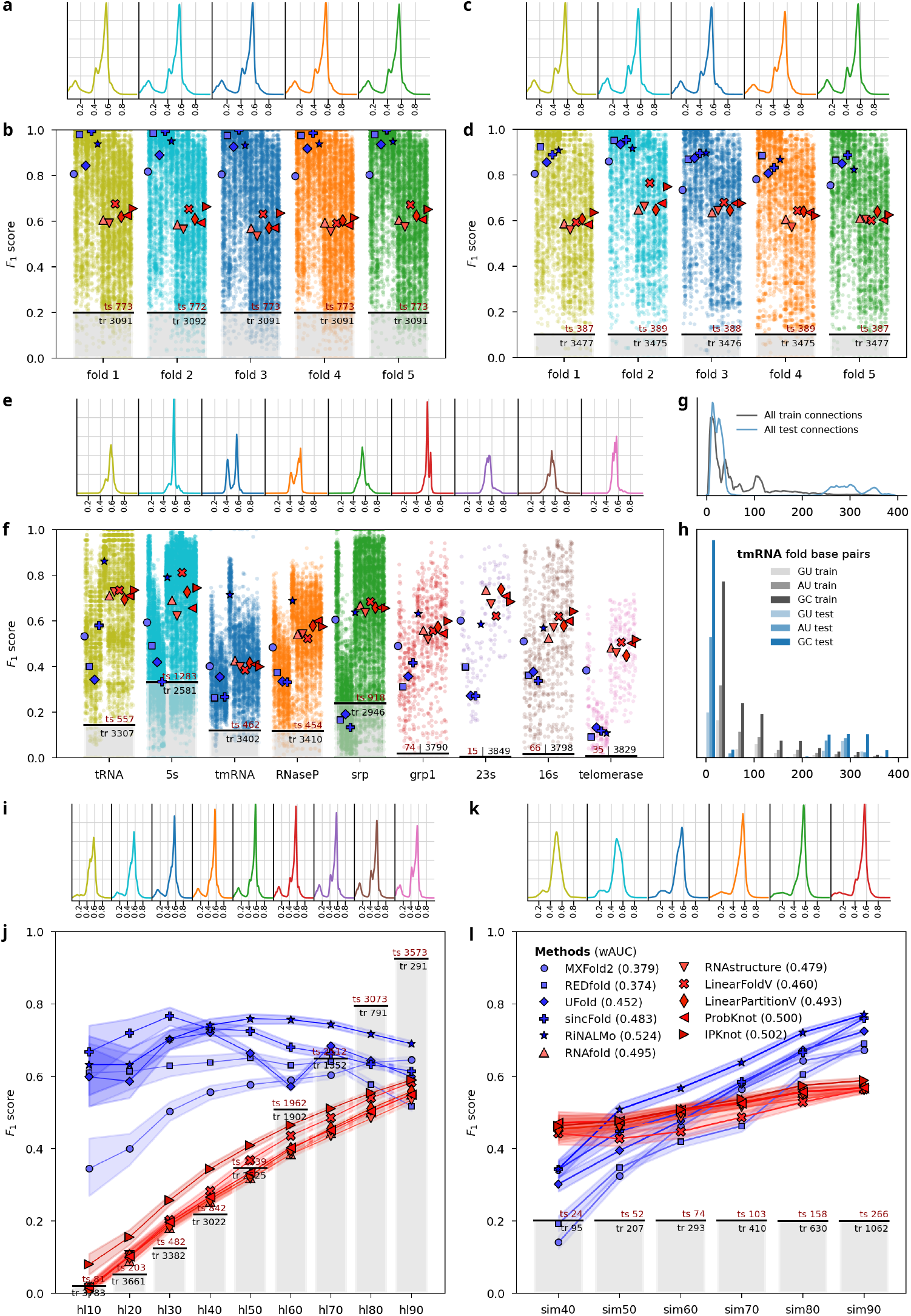
Performance comparison for RNA secondary structure prediction methods according to different cross-validation strategies. k-fold: **a** Distribution of test-to-train structural distances of each fold. **b** Median *F*_1_ for each prediction method indicated with a different marker (DL prediction methods are depicted in blue, classical prediction methods are reported in red). *F*_1_ for each tested sequence is scattered in the background. Gray bars show the proportion of test to train sequences in each fold. At each fold, the test (ts) and train (tr) number of sequences is reported in the bars. **cluster-fold: c** Test-to-train distance distributions. **d** Structure prediction performance. **fam-fold e**: Test-to-train distance distributions. **f** Structure prediction performance. Detailed analysis of connection distances for tmRNA family used in test set of fam-fold strategy. **g** Distribution of distances between bases of canonical pairs in the reference structures of training (gray) and testing (blue) partitions. **h** Distribution of connection distances distribution for canonical base pairs GU, AU and GC. **hl-fold: i** Test-to-train distance distributions. **j** Median *F*_1_ performance for each prediction method, with the 95% confidence interval shaded behind each trend line. **sim-fold: k** Test-to-train distance distributions. **l** Structure prediction performance. wAUC: area under the performance curve for each method, weighted by the difficulty of the partition. Statistical analysis of results in Fig. S1.

### Random and clustering folds

The distributions of test-to-train structural distances for random *k*-fold (Fig. 2.a) and cluster-fold (Fig. 2.c) are nearly identical, even though cluster-fold is intended to better control similarities between partitions. Most sequences have intermediate structural test-to-train distances (peaking at 0.56), but several also have distances below 0.20, which can artificially inflate test results. Regarding prediction performance (Fig. 2.b and Fig. 2.d), both random *k*-fold and cluster-fold show that the average performance of DL models (blue markers) consistently outperforms classical models (red markers), with MXFold2 exhibiting an intermediate performance between both types of methods. This can be also seen in the corresponding statistical analysis with the Critical Difference (CD) diagram (Fig. S1). This indicates that, despite cluster-fold being intended to be a better splitting strategy than random *k*-fold, overestimation of DL models performance still happens in practice.

A stratified per-RNA family variant of random *k*-fold was also evaluated (Fig. S2), showing a performance overestimation for DL methods similar to the obtained with standard *k*-fold. Moreover, higher performance of DL methods can be observed due to increased data-leakage, because stratification ensures that examples from each family are present in both train and test partitions. Regarding cluster-fold, a multi-threshold ranging 30-95% with 5 train/test partitions was also evaluated (Fig. S3.a). Results show that, in spite of using several thresholds, results did not vary substantially, not even in very low similarities. This can be explained by the fact that CD-HIT satisfies intra-cluster similarity but does not control the inter-cluster similarity, thus similar sequences can end up in different clusters, which are then randomly assigned to train or test partitions. Complementary, a multi-threshold cluster-fold at representative level was also evaluated. In this case, only the representatives of clusters were retained to build the train/test partitions, making the problem harder since it leaves just one sequence per cluster. Results show that at low similarity levels performance slightly decreases for DL while remaining stable for classical methods (Fig. S3.b). Below 65% similarity, the *F*_1_ score of all methods fluctuates but it is not largely affected. This can be explained, as in the previous case, by residual data-leakage: although selecting representatives reduces the probability of leakage, it cannot be guaranteed to be entirely eliminated, since the CD-HIT greedy algorithm does not explicitly control inter-cluster similarity (Fig. S3.c-d). Nevertheless, it should be noted that despite these marginal improvements, multi-threshold cluster-fold are not used in RNA secondary structure prediction practice.

### Family folds

The fam-fold test-to-train structural distributions and prediction performances are shown in Fig. 2.e-f. A large heterogeneity can be seen in test-to-train structural distance distributions, where some folds have bimodal distributions and other ones have just a central peak. It is evident that the distances from testing partitions to their corresponding training sets are completely different among families, and most of the folds have distances above 0.4, with some families with peaks in distances beyond 0.60. Instead, 5s and tRNA also have a peak in low distances that might facilitate the prediction of a subgroup of sequences. Overall, there is a large unequal distribution of testing to training distances at every fold. The behavior of the prediction methods in fam-fold (Fig. 2.f) is totally opposite in comparison to random *k*-fold and cluster-fold. With this splitting strategy there is a clear superiority of classical methods over all DL models, in all families, and no matter the proportion of train-to-test samples, with the exception of RiNALMo that achieves results competitive with classical methods in most families. Aggregated over all test sequences, the corresponding CD diagram (Fig. S1) shows that RiNALMo achieves significantly better results than all classical and DL methods, although this global result is not uniform across families. This superior performance over the other DL methods can be attributed to its self-supervised pre-training, where many test sequences available in the public RNA datasets might have already been seen. Also MXFold2 has a particular behavior, with a performance that is higher than DL methods and closer to the classical ones. For example, in the case of srp families, with a lot of testing cases but consequently few train sequences, DL models cannot properly generalize, except for MXFold2, which is actually a hybrid method, and RiNALMo that has been pre-trained. The 5s family also has relatively few sequences for training, but the performances of UFold and REDfold fall less than in other families since these methods overfit to the group of sequences in the peak of smaller test-to-train structural distance. Overall, classical methods provide a moderate behaviour between *F*_1_ = 0.50 and *F*_1_ = 0.70. In 7 out of 9 families, the performance of classical methods and RiNALMo is around *F*_1_ = 0.60 *−* 0.70. In the case of tRNA, RiNALMo performance provides a performance higher than *F*_1_ = 0.80. However, DL models here cannot reach a performance above *F*_1_ = 0.60 for any family. Those are particularly bad in the case of the telomerase RNA family, the longest and the one with the test-to-train structural distance peak at highest distance, where the performance of classical methods is around *F*_1_ = 0.50 but DL models and RiNALMo are in *F*_1_ *≈* 0.10.

We have performed a deeper analysis inside each fold of the fam-fold taking into account the distribution of connection distances for canonical connections, both in train and test partitions. This splitting approach tries to avoid overestimating performance, which could achieve high performance by simple memorization. However, if there are connection patterns that are typical of a family, the fundamental mechanism on which these methods rely (learning from data) is disabled. An example to prove this is shown in Fig. 2.g-h, with the distribution of connection distance between canonical base-pairs (GC, AU and GU) in the training (in shades of blue) and testing (in shades of red) partitions for the tmRNA fold. DL models have a low performance for this family, and it is also the worst predicted by the classical ones. The training partition (with all families except tmRNA) shows a large number of the 3 types of canonical connections at a very short distance, a large peak between 5-10 nt, a peak at 50 nt, and another peak at 100 nt of connection distance. However, the corresponding testing partition (tmRNA family) has a completely different bimodal distribution, with a peak of short-distance connections but a very large number of connections at distances between 250 and 400 nt. Critically, this pattern of connections at such long distances has never been seen during training, making it almost impossible for any ML model to predict them. While this specific example is sufficient to demonstrate the impossibility of learning, similar unmatched patterns of connection distances can also be observed in other families. To quantify these connection distance distributions, we have calculated the Jensen-Shannon distance^50^, for each fold of the fam-fold, between test and train distributions of canonical pairs in the structures (Table S1). It can be observed that the largest distance between test and train distributions are for the tRNA, 5s, tmRNA and telomerase families. Furthermore, in the same way as these simple patterns in canonical connection distances, other more complex patterns specific to a single family could also be found, extending the evidence on the impossibility of learning from the data left in the training set.

### Human learned folds

In the case of hl-fold and sim-fold (Fig. 2.i and 2.k), it can be seen that the distribution of structural distances (top) is similar for folds close to each other. The main difference here with respect to random fold is that each fold has a specific complexity that allows testing prediction models with a known difficulty. Moreover, a continuous gradient of difficulty can be observed in both cases, with more spread distributions in hl-fold, and peaks moving from lower to higher distances in sim-fold. For example, all random folds have a mix of structural complexities, where the low-distance sequences are easy to predict and contribute to inflating the test results. In contrast, while there is a wide variety of different structural distances in the distributions for hl10 and hl90, the difficulty for testing is clearly the opposite. Similarly, in the case of sim-fold, a clear gradualness in the difficulties is also observed. Furthermore, in contrast to all previous cases, the distributions are no longer bimodal but have a single peak, indicating the specific difficulty of the split.

For the performance comparison of prediction methods at the hl-fold (Fig. 2.j), a line plot is used because there is a gradual growing relationship between the splits. As it can be expected, a clear trend can be observed for the performance of classical methods, increasing almost linearly from hl10 to hl90. These models have an extremely low performance in the lowest hl-folds, with median *F*_1_ from 0.0 to 0.20 for hl10 and *F*_1_ from 0.15 to 0.35 in hl20. As the HL threshold increases, the test sets are easier, that is, the number of test sequences for which some human learning has improved over the years increases as well, the HL models improve up to a maximum of average *F*_1_ near 0.60 in hl90. Thus, this strategy works for classic methods as the equivalent of fam-fold for DL models. Differently, DL models have a more stable performance no matter the split in hl-fold. In general, it can be seen that in this splitting strategy DL models provide a more stable and high performance, with *F*_1_ mostly between 0.55 and 0.75. Notably, MXFold2, which is a hybrid method, certainly improves thermodynamic methods performance by almost 20%, yet it falls below most DL methods across almost all folds. UFold has a more variable performance but is always high, with *F*_1_ around 0.70 in the hardest splits of hl-fold. After hl40, RiNALMo is always above all methods, reaching 0.70 at hl90, with overall statistical significance confirmed in the CD diagram (Fig. S1). Intriguingly, DL models increase or maintain performance but only up to a maximum at hl30, after which there are no significant improvements, and then slightly worsen up to hl80. It can be seen that from hl20, a peak starts to appear at low distances in the test to train distances distributions (Fig. 2.i), which increases from hl30, hl40 and on, but it is not at hl10. This peak at low distances indicates a set of sequences that are easy to classify, which may explain the *F*_1_ increase from hl10 to hl30, where data reduction in train has not yet affected model training. As the gray bars indicate, at hl10 there are 5% of sequences in test and 95% in train, while in hl90 is completely the opposite, that is, 90% of sequences in test and 10% in train. This data reduction seems to affect DL methods as the hl-fold increases. To verify its impact, we trained a DL method with the same number of training samples at each hl-fold. Results confirm that, given an equal amount of training data across all folds, the difficulty of learning the sequences decreases as the HL increases (Fig. S4). Therefore, the DL performance peak that was observed at Fig. 2.j occurs because, initially, performance benefits from the reduced difficulty, but then subsequently it begins to be affected by the reduction of training data. Interestingly, at hl90 both HL and ML models converge to a very similar performance, around 0.65. This fact suggests that these test sequences are equally challenging for both types of models.

A deeper analysis of the hl-folds was performed regarding RNA families composition and its relation with classical method performance (Table S2). The same lineal trend in performance in Fig. 2.j is observed for all families. As expected, below hl40, no matter the family proportion in the fold, the classical method has a poor performance. From hl60 and on, the classical method performance reaches a plateau around 0.50-0-60. For the srp family, the classical method provides the worst performance at hl30, hl40 and hl50, slightly improving predictions for the remaining folds up to 0.55. At hl80 and hl90, the best families predicted are 23s and tRNA. For the tmRNA family, similar performance than for tRNA and telomerase at hl40 is achieved, but then performance in tmRNA remains almost unchanged at hl90, achieving the lowest performance in comparison to the other families. For families that would be impossible to predict for a DL method in folds with almost the complete family in the test set (i.e. tmRNA, grp1, 23s, 16s and telomerase), the classical method is capable of providing a prediction with performance 0.40-0.50. Regarding DL method performance in hl-fold, this analysis revealed that when there is a large percentage of a family in the test partition, no matter the fold, the DL method performance is low. On the contrary, as expected, with many samples in training DL methods largely exceed classical ones. Towards the left, where the majority of sequences are in the training set, the few members in the test set tend to be the most difficult, causing performance to drop; towards the right, where the majority are in the test set, there is less information available to learn from. However, there are some families that are especially difficult. In spite of having a large percentage of training samples, DL method performance is low for grp1 and 16s in hl10 and hl20, and RNaseP in hl10. Other families such as tRNA and 5s, are easier to predict even with small training samples (hl90).

Other characterizations of hl-folds regarding number of training/testing samples, mean, maximum and minimum training and testing sequence length, GC content, test-to-train similarity distributions, among others, are provided in Table S3. Recalling that classical methods, by construction, have a linear growth in performance across hl-folds, it is interesting to note that this trend is consistent with the decrease in test GC content (from 59.45% at hl10 to 52.33% at hl90): sequences in lower hl-folds tend to have higher GC content, which increases the number of competing stable alternative structures and makes thermodynamic prediction inherently more challenging. Table S3 also shows that for DL methods, the number of train and test samples at each hl-fold has a high correlation with the *F*_1_ score, as it could be expected for a DL method, since for the largest number of training samples, largest *F*_1_ are achieved. Interestingly, we found that the GC content in train sequences has a high inverse correlation with performance, but GC content in test sequences is not correlated with *F*_1_. Moreover, the absolute difference between the mean training and testing sequence lengths provides one of the strongest correlations with the DL method *F*_1_ across hl-folds. Notably, this difference is what matters rather than the absolute length, since the mean test length alone is uncorrelated with performance. Consistently, the absolute difference between mean lengths reaches its minimum at hl30, precisely the fold at which DL performance peaks. The Jensen-Shannon distance between test and train distributions of connection distances showed a comparably large (inverse) correlation with *F*_1_ score across hl-folds. This is related to the fact that there are connections at distances that are present in the testing partition but absent in the training one, thus negatively affecting the performance of any machine learning method.

### Similarity folds

The performances measured under sim-fold strategy (Fig. 2.l) constitute a more realistic and informative evaluation of models. It shows how the model will behave in future situations where test sequences present different levels of homology with the training set. In the case of classical methods, it can be seen that the average performance for most similarity levels is around *F*_1_ = 0.50 *−* 0.55. Even in the cases with the highest similarity between train and test (sim90), the best performance is below 0.60. This is expected because the remaining data in the training sets are not used to re-tune the parameters of classical methods. However, at the hardest sim40 split, it seems that the selected test examples are also different from the sequences that were used to build the thermodynamics models, and thus performance is the lowest. Interestingly, these sequences in the low similarity splits are not only very different from each other, being a challenge for learning-based methods, but also fold according to laws that defy the known folding rules and the measured thermodynamic parameters with which classical methods were built and tuned for decades. From sim50 onward, DL models are capable of providing a minimal prediction performance, even though the training sequences were so different and there are just a few examples for training. In other words, these splits present a double difficulty for DL models, due to the difference between the training and testing patterns, and due to the few sequences available for training. Notably, these folds are as hard as the hardest fam-fold splits for DL models, but now being equally difficult for classical methods as well. As the similarity between train and test increases, DL models take advantage of this fact and improve almost linearly up to 0.80 clearly surpassing classical methods by a 20% at the highest and easiest split (sim90). A detailed analysis of per-sequence test performance of classical and DL methods as a function of the maximum similarity to the training set at sim90 reveals that the performance of classical methods remains almost unaltered no matter the similarity (Fig. S5). On the contrary, and as it could be expected from DL methods, performance is indeed correlated with test-to-train similarity. Remarkably, RiNALMo outperforms all DL methods along all folds, presumably due to its pre-training; only sincFold matches its performance at sim40 and sim90. Additionally for each method, the weighted area under the curve (wAUC) is indicated in the figure legend, providing a global measure of performance for each method, weighted by the difficulty of the partition. That is, the best performances for high similarities (e.g., sim90) will have less weight (0.10) in the average, thus giving more relevance to performances in the most difficult cases (see Methods for more details). Among the evaluated models, IPknot achieved the highest overall score among classical methods with wAUC 0.502. Within the DL models, RiNALMo achieved the best wAUC (0.524), followed by sincFold (0.483) and UFold (0.452). In contrast, other DL approaches such as MXFold2 and REDfold have lower performances, failing below 0.380. In this scenario, MXFold2 behaves more like a DL model than a classical one, clearly surpassing classical methods at sim80 and sim90, but showing poor performance at low similarity thresholds. In the overall assessment, the wAUC provided a comprehensive measure of how each type of method performs across the whole range of similarities, revealing a much smaller gap between them than either random k-fold or fam-fold would suggest, as these exaggerate it in opposite directions.

### Complementarity between methods and validation on a second dataset

For all the strategies evaluated in this study, the per-sequence correlation between *F*_1_ scores of all folding methods was calculated. Results show that for both random *k*-fold and cluster-fold, DL-based methods provide highly similar predictions among them, and the same happens with classical methods (Fig. S6). These matrices reveal two clearly defined blocks, suggesting that both types of methods have complementary strengths. MXFold2, as a hybrid method, shows an intermediate behavior with moderate correlations to both groups. In the fam-fold scenario, correlations highly depend on the RNA family. For example, for grp1 and 16s, DL and classical methods provide similar predictions, while for the 23s family, performances of classical methods are even negatively correlated with sincFold and UFold, emphasizing the complementary behaviour. For tmRNA, tRNA, 5s and telomerase, while classical methods are fairly consistent among them, each DL method provides different predictions (i.e. without a defined block in this type of methods). Notably, the clear two-block structure observed under *k*-fold and cluster-fold disappears under fam-fold, and intra-DL correlations become more heterogeneous. This suggests that when homology between train and test is removed, DL methods no longer fail consistently on the same sequences. However, it should be noted that for some families, overall performance levels are so low for all methods, therefore per-sequence correlations should be interpreted with caution. In the hl-fold, no matter the fold, DL methods provide highly correlated predictions. However, classical methods at low hl-folds show little correlation both among them and with DL methods. Interestingly, from hl40 onwards, LinearFoldV and IPknot show higher correlations with DL methods than with classical ones. As expected, the two-blocks pattern is rising when increasing the HL threshold up to hl90. At this point, the correlation pattern closely resembles that of *k*-fold. For sim-folds strategy, classical methods show very high correlations at sim40, while DL methods are moderately correlated, with the exception of sincFold and RiNALMo which show a low correlation (0.35). Notably, MXFold2 presents predictions that are very different from all other types of methods, showing no correlation with classical methods and low correlations with DL methods as well, suggesting that under strict similarity constraints its hybrid nature leads to a complementary behavior in predictions. From sim50 onwards, a gradual convergence toward the two-block structure is observed.

All cross-validation strategies and folding methods have been applied also to the bpRNA dataset^51^ (Fig. S7). At random *k*-fold and cluster-fold, all DL methods have better performance than classical methods, however with lower values than in ArchiveII. Classical methods maintain a moderate performance. As stated for the other dataset, there is also no significant difference between random *k*-fold and cluster-fold strategies. At fam-fold the behavior is also similar to the one observed: classical methods are superior to DL ones, with some specific RNA families equally challenging for both types. At hl-fold the performance of all methods is low but increasing as the folds become easier, with DL methods also increasing and slightly above classical ones. Also here, at hl90 both HL and DL models converge to a very similar performance, in this case around 0.40 *−* 0.50. In sim-fold, classical methods are not affected while DL performance improves as similarity between train and test increases. In this dataset, the performance of classical methods is more stable around 0.50 no matter the similarity, and thus they obtain the highest wAUC values. At sim40 and sim50, classical methods provide higher performance as in the other dataset; while at sim90 the DL models sincFold and UFold provide the best *F*_1_. Notably, RiNALMo does not retain here the advantage it showed in ArchiveII, with a wAUC below sincFold. This is in line with our interpretation that its advantage comes from pre-training exposure, which cannot be assumed for the more diverse set of families in bpRNA. In general, the previous analysis of all cross-validation strategies holds on this dataset, confirming that sim-fold, together with the wAUC metric, provides the most informative assessment of RNA secondary structure prediction methods performance across the full range of similarities.

## Concluding remarks

In this study we analyzed in detail the most commonly used cross-validation strategies for RNA secondary structure prediction. We have also explored two novel evaluation methodologies: one designed to test in a hard way both ML as well as HL models; and another one to explore the full range of sequence similarities from test to train partitions. All validation strategies were applied to state-of-the-art prediction methods, including both classical thermodynamics approaches and the newest DL-based ones, on two datasets differing in family composition and degree of homology. The provided experimental support showed that in spite of enabling some degree of homology control, cluster-fold provides the same limited information about generalization capabilities as random *k*-fold, which also holds for its multi-threshold and cluster-representative variants. A detailed analysis of the train and test partitions showed that the family fold splitting violates the main hypothesis behind ML models, because key patterns in test families are not present in the training partition, making learning a theoretically impossible task in this scenario.

The comprehensive analysis of the commonly used splitting strategies reveals that the distance distribution from testing to training partition introduces biases that favor some methods over others. The low distances in *k*-fold and cluster-fold give undue advantage to DL models, whereas the opposite is the case for the high distances of fam-fold. Consequently, none of these three splitting strategies provide a fair comparison of generalization capabilities. While random *k*-fold and cluster-fold clearly benefit DL models due to data leakage from uncontrolled similarity, fam-fold suffers from a different kind of leakage: the classical methods benefit from the experimental measurements and knowledge used for their original development and parameter tuning, which may already include the families being tested. A comparable caveat applies to pre-trained models: their exposure to public RNA data precedes, and is unaffected by, any partitioning of the dataset.

With the hl-fold and sim-fold strategies, we recreated fair and realistic evaluation scenarios, showing how each predictor is expected to behave in real-world scenarios where different levels of homologies will be received. These strategies for cross-validation provide a wide spectrum of challenges for performance evaluation at all levels of difficulty and for all types of methods, ensuring a balanced benchmark for the field. The sim-fold strategy, which is based on direct pairwise sequence similarities, strictly ensures, by design, that there is no data leakage between test and train partitions. Unlike the commonly used strategies, sim-fold characterizes performance of RNA secondary structure prediction methods across the full range of test-to-train similarities while strictly controlling data leakage at each level.

## Materials and methods

### Data and cross-validation setup

In this study we use the ArchiveII dataset^52^, which is the most commonly employed gold standard for evaluating the performance of RNA secondary structure prediction methods. ArchiveII contains 3,975 RNA structures from 9 well-known RNA families: 5S rRNAs, SRP RNA, tRNA, tmRNA, RNase P RNA, 16S rRNA, grp1, Telomerase RNA and 23S rRNA. As suggested in^16^, sequences longer than 512 nucleotides were filtered, resulting in a final dataset of 3,864 sequences.

The bpRNA dataset is also used, as proposed in^24^, with annotated secondary structures from bpRNA^51^. This dataset has 13,419 RNAs, after excluding sequences with *>*80% of identity with CD-HIT and longer than 500 nucleotides. We have also included the bpRNA-new dataset^53^, which contains 5,401 sequences from new RNA families different from the bpRNA dataset. Both sets were aggregated and sequences with ambiguous nucleotides were removed. We used the resulting 18,537 sequences to build the random *k*-fold, cluster-fold, hl-fold and sim-fold partitions, while for fam-fold, bpRNA was used for training and bpRNA-new for testing.

For the random *k*-fold strategy, we adopted the original *k* = 5 configuration from ArchiveII for all the methods evaluated. For assessing cross-family generalization, fam-fold cross-validation was performed by leaving one family out for testing per fold, and the remaining families were used for training. In the case of bpRNA, the 20 largest RNA families were analysed. For cluster-fold splitting, CD-HIT program was used with a threshold of 80% as in previous works^24, 32^. Then, the clusters of sequences were randomly assigned to each partition. In the hl-fold strategy, hl*τ* splits were defined by assigning to the training set all the sequences for which a consensus of classical methods (RNAfold, RNAstructure, LinearFold and LinearPartition) provide a performance higher than the given threshold *τ*, leaving the remaining sequences for testing. This way, several hl-folds were obtained, from hl10 to hl90. Finally, in the sim-fold strategy, pairwise sequence similarity for each dataset was calculated with the pairwise2 module from Biopython, which uses the Smith-Waterman local alignment algorithm with gaps penalization, with similarity defined as the number of identical positions divided by the length of the longer sequence. Multiple splits were created with similarities ranging from 10 to 90. Within splits there are no redundant sequences because the maximum sequence similarity is guaranteed in this splitting strategy. Bootstrapping was performed within each split in order to obtain 10 folds, with the corresponding train and test partitions. The sim10, sim20, and sim30 splits were not used due to an insufficient number of sequences fulfilling the similarity conditions.

### Performance and distance measures

In this study the focus of performance evaluation is on the predicted base pairs, in comparison to a reference (ground truth) structure. Pairs present in both the prediction and in the reference structure are the true positives (TP), while pairs predicted but not present in the reference structure are false positives (FP). Similarly, a pair in the reference structure that is not predicted is a false negative (FN). To fully characterize the successes and failures of structure prediction, the F1 score is defined as

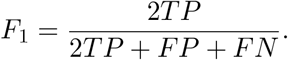

For each strategy evaluated and method performance obtained, the corresponding statistical significance analysis is provided through critical difference (CD) diagrams^54^.

For each method evaluated in the sim-fold strategy, a weighted area under the performance curve (wAUC) is calculated as

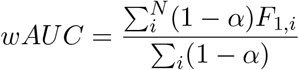

where *N* is the number of splits and, for each split *i*, the *F*_1_ of the method is weighted by the difficulty factor 1 *− α*, with *α* = *σ/*100.

### RNA secondary structure prediction methods

The classical methods evaluated in this study are based on thermodynamic principles, primarily utilizing the Turner nearest-neighbor parameters^55^ to evaluate energy changes in RNA loops and helices.

**RNAfold** (v2.5.0)^47^ is the core program of the ViennaRNA package and the most widely used algorithm for RNA secondary structure prediction. It predicts secondary structure by finding the minimum free energy (MFE) conformation using dynamic programming^48^, and the partition function^56^ to determine base-pairing probabilities.

**RNAstructure** (v6.4)^57^, similarly to RNAfold, works by calculating the most stable MFE folding of a sequence via dynamic programming, with ability to incorporate experimental chemical probing data.

**ProbKnot** (v6.4)^58^ predicts base-pair probabilities using a partition function that excludes pseudoknots, and then assembles the maximum expected accuracy (MEA) structure from those probabilities. As a post-processing step, the algorithm removes unstable helices.

**IPknot** (v1.0.0)^59^ is specifically designed to predict pseudoknoted structures. It decomposes a structure into a set of pseudoknot-free substructures, approximating their base-pairing probability distributions, and solves the MEA maximization using integer programming instead of standard dynamic programming.

**LinearFold**^10^ is an approximate algorithm that adapts the MFE framework to achieve a linear runtime. It processes the sequence from left to right, utilizing a heuristic beam search to prune the conformational search space during dynamic programming.

**LinearPartition**^60^ is the linear-time counterpart for partition function calculation, approximating base-pairing probabilities. It employs a similar beam pruning heuristic to retain only the most promising candidates derived from the Turner model, reducing computational overhead.

**MXFold2** (v0.1.1)^53^ is a hybrid model that integrates the Turner thermodynamic parameters with folding scores from a DL model. The DL model is trained to minimize the structured hinge loss function with thermodynamic regularization. The final model returns the secondary structure that maximizes the combined thermodynamic and learned scores through Zuker dynamic programming.

The DL and hybrid models for this work were selected based on their availability as open access models that can be trained and tested with custom datasets from scratch. Therefore^24, 61–65^ could not be included in this work. The following models were all re-trained from scratch in each corresponding dataset with each cross validation strategy.

**UFold** (v1.2)^32^ is a DL model that represents the RNA sequence through all the possible 16 basepairing maps. These input matrices are treated as an image with 16 channels, plus a channel for the pairing probability between each base pair. A standard U-Net encoder-decoder framework is used to predict the contact score map between all nucleotides. Following the original source code released by the authors, the model was trained 100 epochs without validation split, selecting the last epoch checkpoint as the final model parameters, with all hyperparameters by default.

**REDfold**^33^ is a DL model that also utilizes a U-Net encoder-decoder to learn dependencies within the RNA sequence. It incorporates symmetric skip connections to propagate activation information across layers, followed by an output post-processing step with constrained optimization. This model was trained with the default configuration, for 200 training epochs without validation set, using for inference the last checkpoint.

**sincFold**^66^ is an end-to-end DL model based on ResNet bottlenecks, designed to capture both short- and long-range dependencies in the RNA sequence. It has a two stage encoding process: a sequence encoding in 1D, to learn local context features, followed by a pairwise encoding in 2D, to capture distant relationships. This model was trained with early stopping criteria with 5 epochs patience and maximum 100 epochs. The best checkpoint was selected using 10% of the train data for validation.

**RiNALMo**^49^ is one of the largest RNA language models, with 650 million parameters, pretrained on 36 million unique ncRNA sequences from several RNA databases. Its architecture is a BERT-style encoder-only, with 33 Transformer blocks having 20 multi-heads attention each, with residual connections and layer normalization to stabilize training, followed by a 2 layer feed-forward network. No modifications were made to the architecture reported in the original paper. The prediction head^30^ was trained for a maximum of 30 epochs, with early stopping using a patience of 5 epochs. The same validation splits used for the other models were employed for RiNALMo, as defined in the corresponding splitting files. No additional hyperparameter tuning was performed, and the model checkpoint achieving the best validation performance was selected as the final model.

## Data availabiity

Data and source code are available at https://github.com/sinc-lab/xvalRNAfolding

## Author contributions statement

D.H.M conceived the research goals; D.H.M, L.B., G.K., L.D.P, and M.G. wrote the code and conducted the experiments; D.H.M. prepared the figures; G.S. wrote the original draft; G.S. and D.H.M. analysed the results and wrote the manuscript; all authors reviewed, edited and approved the manuscript.

## References

1. J Henninger, J Young, An RNA-centric view of transcription and genome organization. Mol. Cell 84, 3627–3643 (2024).

2. H Hwang, et al., Big data and deep learning for RNA biology. Exp. Mol. Medicine 56, 1293–1321 (2024).

3. B Furtig, et al., NMR Spectroscopy of RNA. ChemBioChem 4, 936–962 (2003).

4. J Zhang, Y Fei, L Sun, Q Zhang, Advances and opportunities in rna structure experimental determination and computational modeling. Nat. Methods 19, 1193–1207 (2022).

5. R Townshend, et al., Geometric deep learning of rna structure. Science 373, 1047–1051 (2021).

6. A Mustoe, HA Hashimi, C Brooks, Secondary structure encodes a cooperative tertiary folding funnel in the azoarcus ribozyme. Nucleic Acids Res. 44, 402–412 (2015).

7. D Kwon, Rna function follows form – why is it so hard to predict? Nature 639, 1106–1108 (2025).

8. Y Shu, et al., Advances in rna secondary structure prediction and rna modifications: Methods, data, and applications (2025).

9. I Hofacker, et al., Fast folding and comparison of RNA secondary structures. Monatshefte fur Chemie Chem. Mon. 125, 167–188 (1994).

10. L Huang, et al., LinearFold: linear-time approximate RNA folding by 5’-to-3’ dynamic programming and beam search. Bioinformatics 35, i295–i304 (2019).

11. M Justyna, M Antczak, M Szachniuk, Machine learning for rna 2d structure prediction benchmarked on experimental data. Briefings Bioinforma. 24, 1–10 (2023).

12. J Jumper, et al., Highly accurate protein structure prediction with AlphaFold. Nature 596, 583–589 (2021).

13. R Geirhos, et al., Shortcut learning in deep neural networks. Nat. Mach. Intell. 2, 665–673 (2020).

14. G Sacco, G Bussi, G Sanguinetti, Machine learning for rna secondary structure prediction: a review of current methods and challenges. RNA 1, rna.080840.125 (2026).

15. M Szikszai, et al., Deep learning for rna secondary structure determination: gauging generalizability and broadening the scope of traditional methods. RNA 32, 428–442 (2026).

16. M Szikszai, et al., Deep learning models for RNA secondary structure prediction (probably) do not generalize across families. Bioinformatics 38, 3892–3899 (2022).

17. B Heil, et al., Reproducibility standards for machine learning in the life sciences. Nat. Methods 18, 1132–1135 (2021).

18. I Walsh, et al., DOME: recommendations for supervised machine learning validation in biology. Nat. Methods 18, 1122–1127 (2021).

19. T Hastie, J Friedman, R Tibshirani, The elements of statistical learning: data mining, inference, and prediction. Springer (2009).

20. A Florensa, et al., SpanSeq: similarity-based sequence data splitting method for improved development and assessment of deep learning projects. NAR Genomics Bioinforma. 6, 1–10 (2024).

21. V Vapnik, The Nature of Statistical Learning Theory. Springer (1995).

22. Y Yang, et al., Genome-scale characterization of RNA tertiary structures and their functional impact by RNA solvent accessibility prediction. RNA 23, 14–22 (2016).

23. I Guruge, et al., B-factor profile prediction for RNA flexibility using support vector machines. J. Comput. Chem. 39, 407–411 (2017).

24. J Singh, J Hanson, K Paliwal, Y Zhou, Rna secondary structure prediction using an ensemble of two-dimensional deep neural networks and transfer learning. Nat. Commun. 10, 1–10 (2019).

25. L Fu, et al., CD-HIT: accelerated for clustering the next-generation sequencing data. Bioinformatics 28, 3150–3152 (2012).

26. F Kallenborn, et al., Gpu-accelerated homology search with mmseqs2. Nat. Methods 22, 2024–2027 (2025).

27. W Hogg, S Villar, Position: is machine learning good or bad for the natural sciences? in Proceedings of the 41st International Conference on Machine Learning, ICML’24. (JMLR.org), (2024).

28. A de Lajarte, et al., Diverse database and machine learning model to narrow the generalization gap in RNA structure prediction. bioRxiv (2024).

29. L Bugnon, et al., Secondary structure prediction of long noncoding RNA: review and experimental comparison of existing approaches. Briefings Bioinforma. 23, 1–10 (2022).

30. L Zablocki, et al., Comprehensive benchmarking of large language models for RNA secondary structure prediction. Briefings Bioinforma. 26, 1–10 (2025).

31. U Hobohm, et al., Selection of representative protein data sets. Protein Sci. 1, 409–417 (1992).

32. L Fu, et al., UFold: fast and accurate RNA secondary structure prediction with deep learning. Nucleic Acids Res. 50, e14–e14 (2021).

33. C Chen, Y Chan, REDfold: accurate RNA secondary structure prediction using residual encoder-decoder network. BMC Bioinforma. 24, 1–10 (2023).

34. BS Booy, et al., RNA secondary structure prediction with convolutional neural networks. BMC Bioinforma. 23, 1–10 (2022).

35. Y Wang, et al., Enhanced RNA secondary structure prediction through integrative deep learning and structural context analysis. Nucleic Acids Res. 53, 1–10 (2025).

36. E Rivas, R Lang, S Eddy, A range of complex probabilistic models for RNA secondary structure prediction that includes the nearest-neighbor model and more. RNA 18, 193–212 (2011).

37. M Szikszai, et al., RNA3DB: A structurally-dissimilar dataset split for training and benchmarking deep learning models for rna structure prediction. J. Mol. Biol. 436, 168552 (2024).

38. JA Cruz, et al., <i>rna-puzzles</i> : A casp-like evaluation of rna three-dimensional structure prediction. RNA 18, 610–625 (2012).

39. D Mathews, et al., Incorporating chemical modification constraints into a dynamic programming algorithm for prediction of RNA secondary structure. Proc. Natl. Acad. Sci. 101, 7287–7292 (2004).

40. D Proctor, et al., Isolation and characterization of a family of stable RNA tetraloops with the motif ynmg that participate in tertiary interactions. Biochemistry 41, 12062–12075 (2002).

41. V Antao, et al., A thermodynamic study of unusually stable RNA and DNA hairpins. Nucleic Acids Res. 19, 5901–5905 (1991).

42. V Antao, I Tinoco, Thermodynamic parameters for loop formation in RNA and DNA hairpin tetraloops. Nucleic Acids Res. 20, 819–824 (1992).

43. D Groebe, O Uhlenbeck, Characterization of RNA hairpin loop stability. Nucleic Acids Res. 16, 11725–11735 (1988).

44. C Flamm, et al., Caveats to deep learning approaches to RNA secondary structure prediction. Front. Bioinforma. 2, 1–10 (2022).

45. W Murdoch, et al., Definitions, methods, and applications in interpretable machine learning. Proc. Natl. Acad. Sci. 116, 22071–22080 (2019).

46. S Cranford, One more thing … patterns 2.0. Patterns 4, 100677 (2023).

47. R Lorenz, et al., ViennaRNA package 2.0. Algorithms for Mol. Biol. 6, 1–10 (2011).

48. M Zuker, P Stiegler, Optimal computer folding of large RNA sequences using thermodynamics and auxiliary information. Nucleic Acids Res. 9, 133–148 (1981).

49. RJ Penić, T Vlašić, RG Huber, Y Wan, M Šikić, Rinalmo: general-purpose rna language models can generalize well on structure prediction tasks. Nat. Commun. 16 (2025).

50. CD Manning, H Schütze, Foundations of Statistical Natural Language Processing. (The MIT Press), (1999).

51. P Danaee, et al., bprna: large-scale automated annotation and analysis of rna secondary structure. Nucleic Acids Res. 46, 5381–5394 (2018).

52. M Sloma, D Mathews, Exact calculation of loop formation probability identifies folding motifs in RNA secondary structures. RNA 22, 1808–1818 (2016).

53. K Sato, et al., RNA secondary structure prediction using deep learning with thermodynamic integration. Nat. Commun. 12, 1–10 (2021).

54. J Demšar, Statistical comparisons of classifiers over multiple data sets. J. Mach. Learn. Res. 7, 1–30 (2006).

55. D Turner, D Mathews, NNDB: the nearest neighbor parameter database for predicting stability of nucleic acid secondary structure. Nucleic Acids Res. 38, D280–D282 (2009).

56. J McCaskill, The equilibrium partition function and base pair binding probabilities for RNA secondary structure. Biopolymers 29, 1105–1119 (1990).

57. J Reuter, D Mathews, RNAstructure: software for RNA secondary structure prediction and analysis. BMC Bioinforma. 11, 1–10 (2010).

58. S Bellaousov, D Mathews, ProbKnot: Fast prediction of RNA secondary structure including pseudo-knots. RNA 16, 1870–1880 (2010).

59. K Sato, et al., IPknot: fast and accurate prediction of RNA secondary structures with pseudoknots using integer programming. Bioinformatics 27, i85–i93 (year?).

60. H Zhang, et al., LinearPartition: linear-time approximation of RNA folding partition function and base-pairing probabilities. Bioinformatics 36, i258–i267 (2020).

61. N Calonaci, et al., Machine learning a model for RNA structure prediction. NAR Genomics Bioinforma. 2, 1–10 (2020).

62. H Zhang, et al., A new method of RNA secondary structure prediction based on convolutional neural network and dynamic programming. Front. Genet. 10, 1–10 (2019).

63. L Quan, et al., Developing parallel ant colonies filtered by deep learned constrains for predicting RNA secondary structure with pseudo-knots. Neurocomputing 384, 104–114 (2020).

64. L Wang, et al., DMfold: A novel method to predict rna secondary structure with pseudoknots based on deep learning and improved base pair maximization principle. Front. Genet. 10, 1–10 (2019).

65. H Yonemoto, K Asai, M Hamada, A semi-supervised learning approach for RNA secondary structure prediction. Comput. Biol. Chem. 57, 72–79 (2015).

66. L Bugnon, et al., sincFold: end-to-end learning of short- and long-range interactions in RNA secondary structure. Briefings Bioinforma. 25, bbae271 (2024).

