## Supplementary Material for "Homology-aware cross-validation strategies for generalization assessment in RNA secondary structure prediction"

#### Supplementary figures

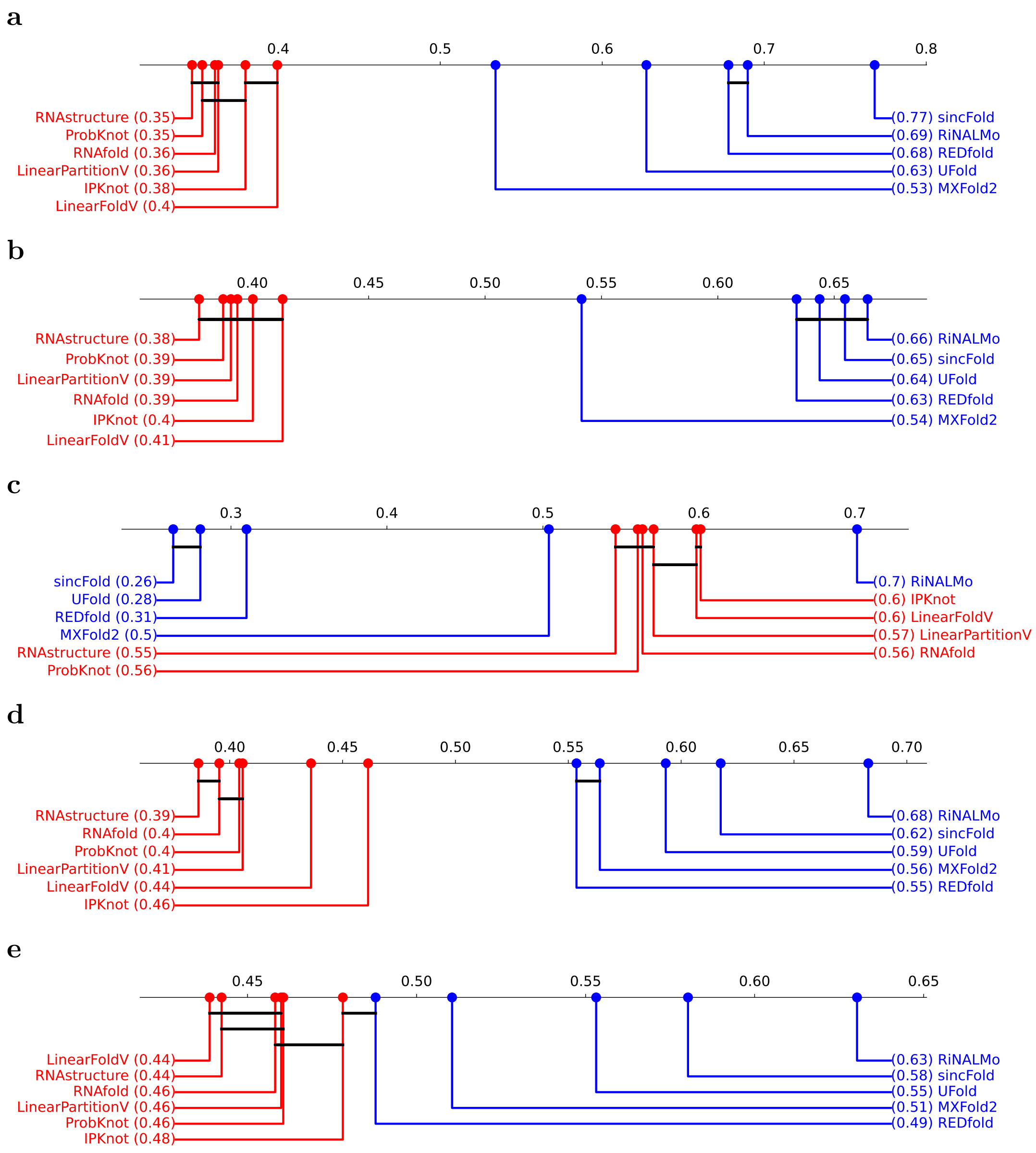


**Fig. S1.** Critical difference diagram of the performance for RNA secondary structure prediction methods according to different cross-validation strategies. **a** k-fold, **b** cluster-fold, **c** family-fold, **d** hl-fold and **e** sim-fold.

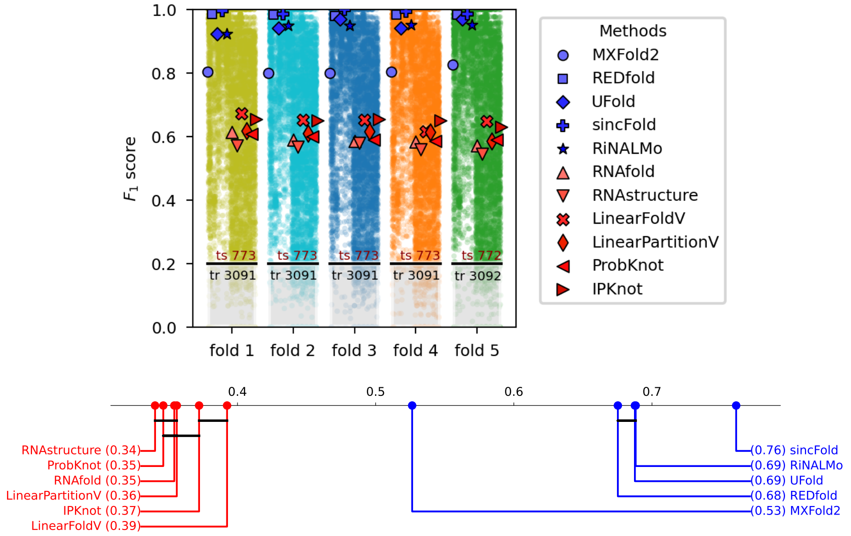


**Fig. S2.** Performance and critical difference diagram of prediction methods for k-fold stratified by families of Archive II.

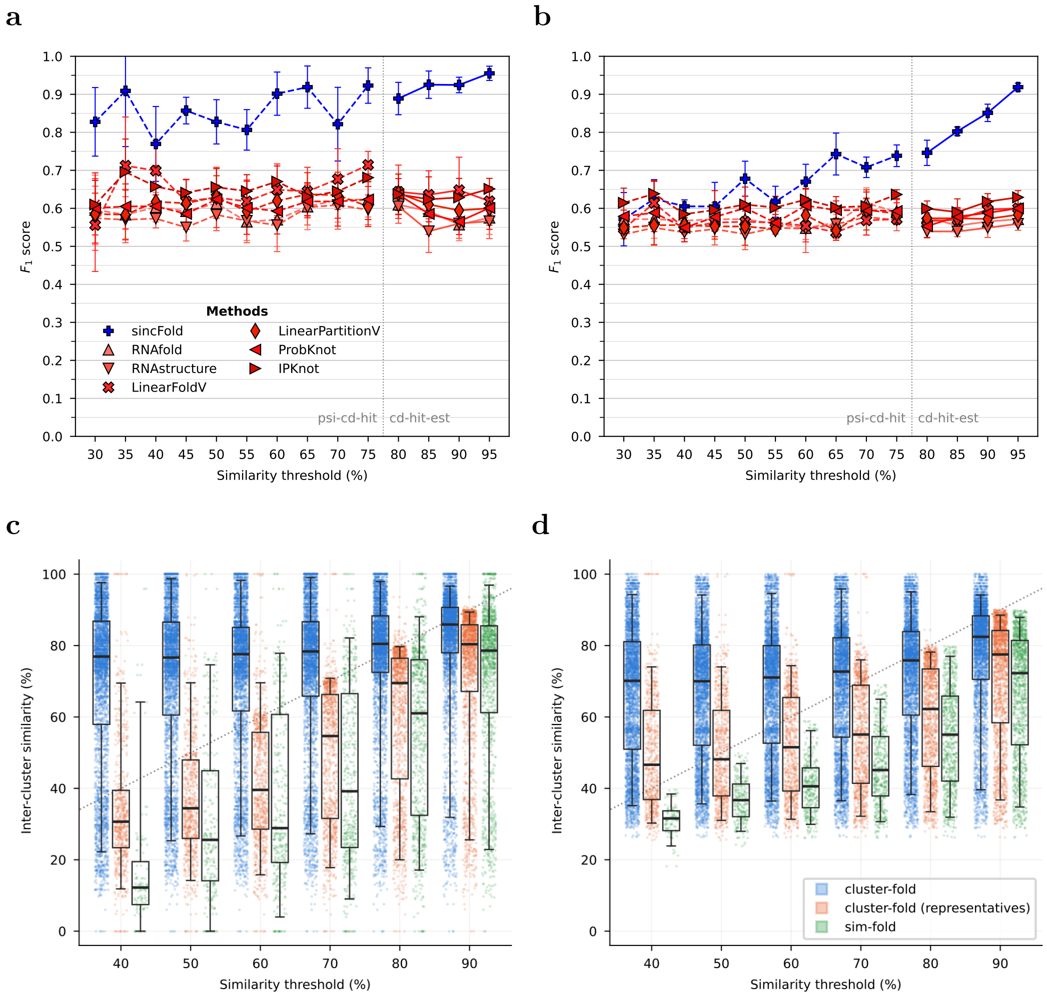


**Fig. S3.** Performance of prediction methods for multi-threshold cluster-fold cross-validation variants with thresholds values ranging 30-95% in the Archive II dataset. **a** Using full clusters to build k-fold partitions for train/test. **b** Using only one single cluster representative (medoid) per cluster to build k-fold train/test partitions. **c** Data leakage measured as BLASTN identity to the nearest sequence in a different cluster. **d** Data leakage measured with the Smith-Waterman local alignment algorithm.

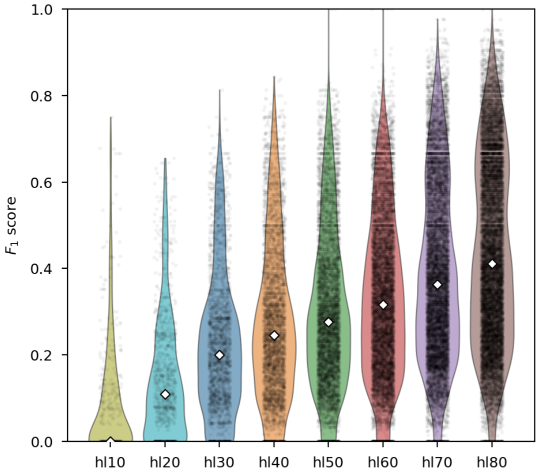


**Fig. S4.** Performance results for sincFold at hl-folds with the same amount of train data in each fold.

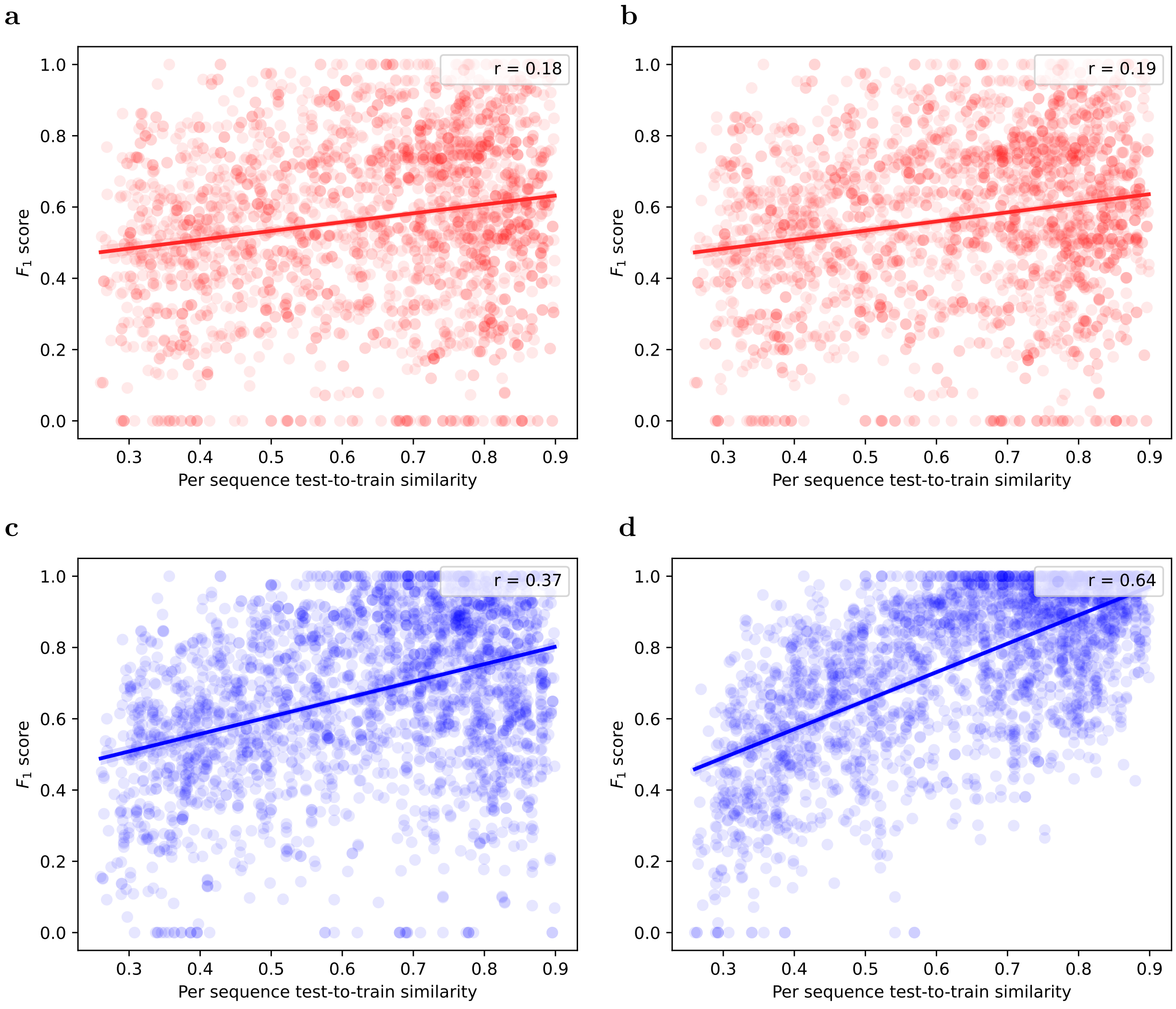


**Fig. S5.** Detailed analysis of per-sequence test $F_1$ score (y-axis) of classical methods (red) and DL methods (blue), as a function of the maximum similarity to the training set (x-axis) at sim90. For each test sequence, its maximum similarity to all the sequences of the train set is depicted. **a** RNAfold. **b** LinearPartitionV. **c** MXFold2. **d** sincFold.

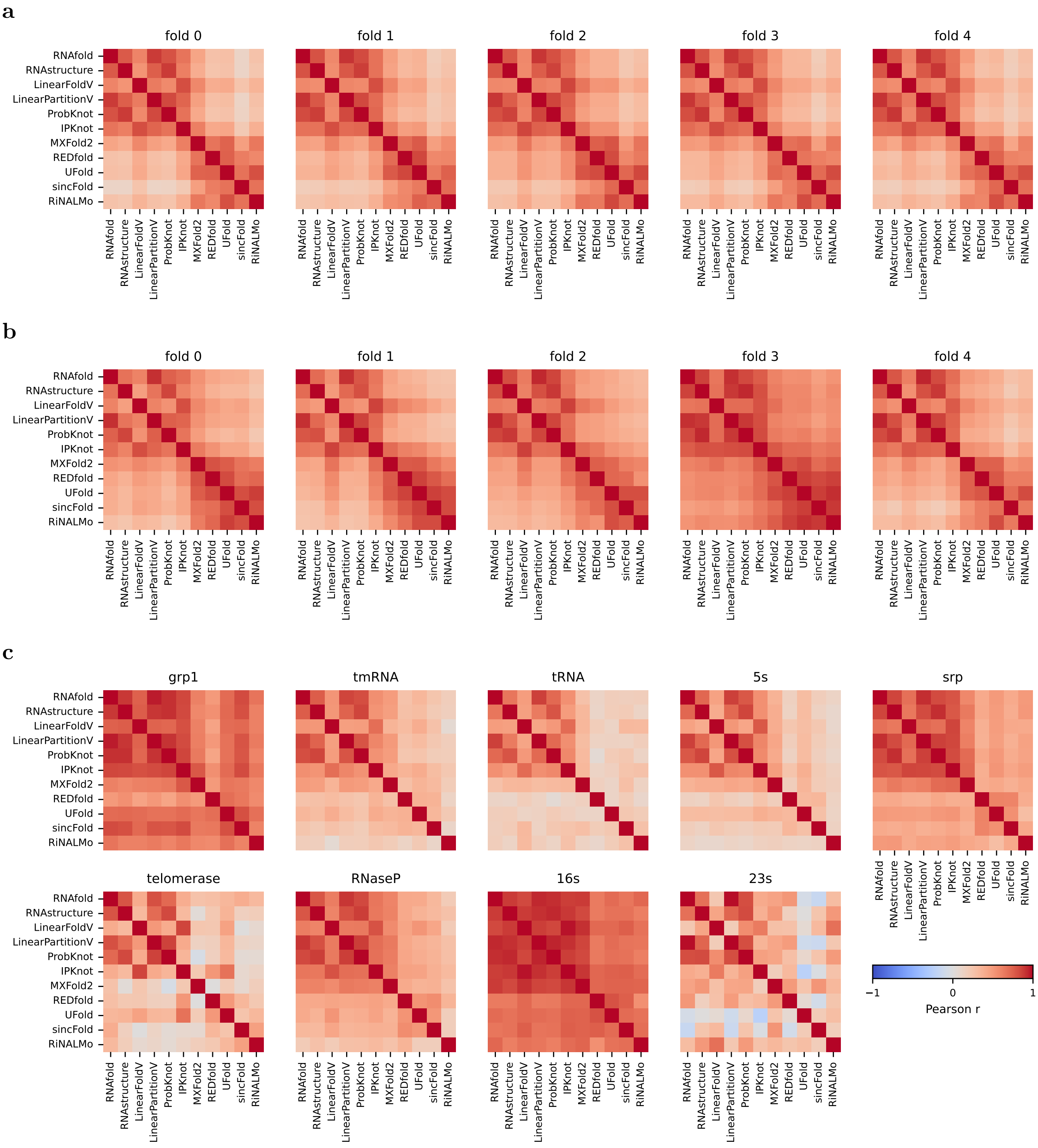
a
fold 0
fold 1
fold 2
fold 3
fold 4
b
fold 0
fold 1
fold 2
fold 3
fold 4
c
grp1
tmRNA
tRNA
5s
srp
telomerase
RNaseP
16s
23s
RNAfold
RNAstructure
LinearFoldV
LinearPartitionV
ProbKnot
IPKnot
MXFold2
REDfold
UFold
sincFold
RiNALMo
−1
0
1
Pearson r

**d**

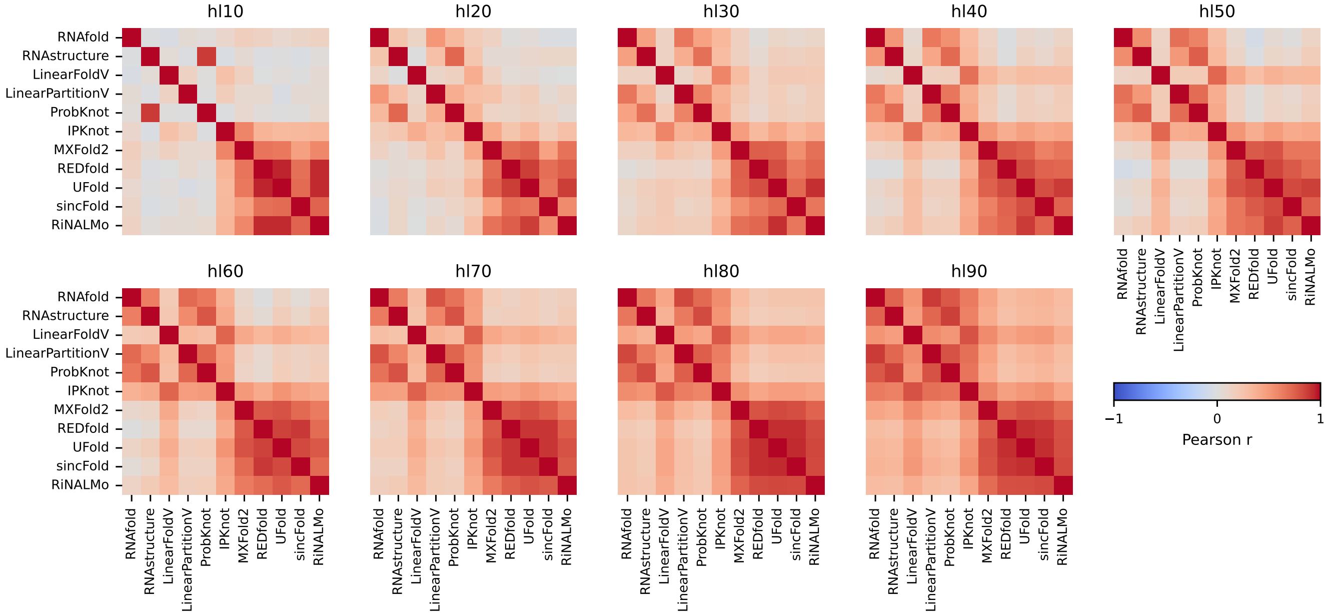


**e**

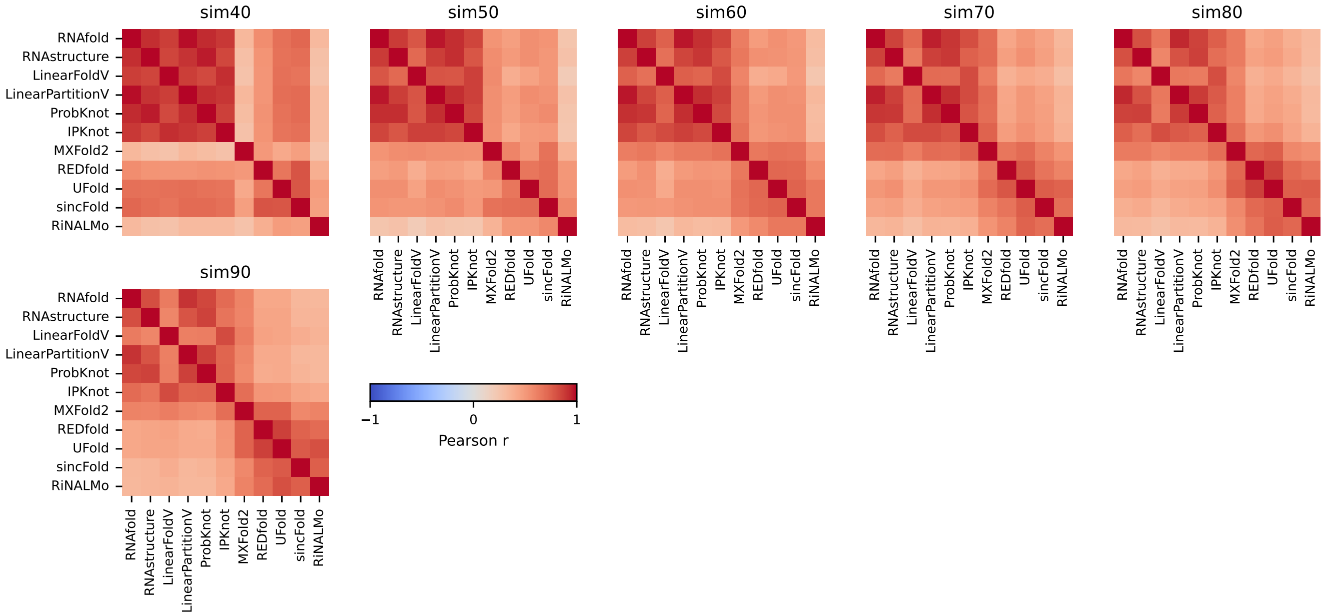


**Fig. S6.** Per-sequence correlations between $F_1$ scores of DL-based and classical methods, for each cross-validation strategy. **a** k-fold. **b** cluster-fold. **c** fam-fold. **d** hl-fold. **e** sim-fold.

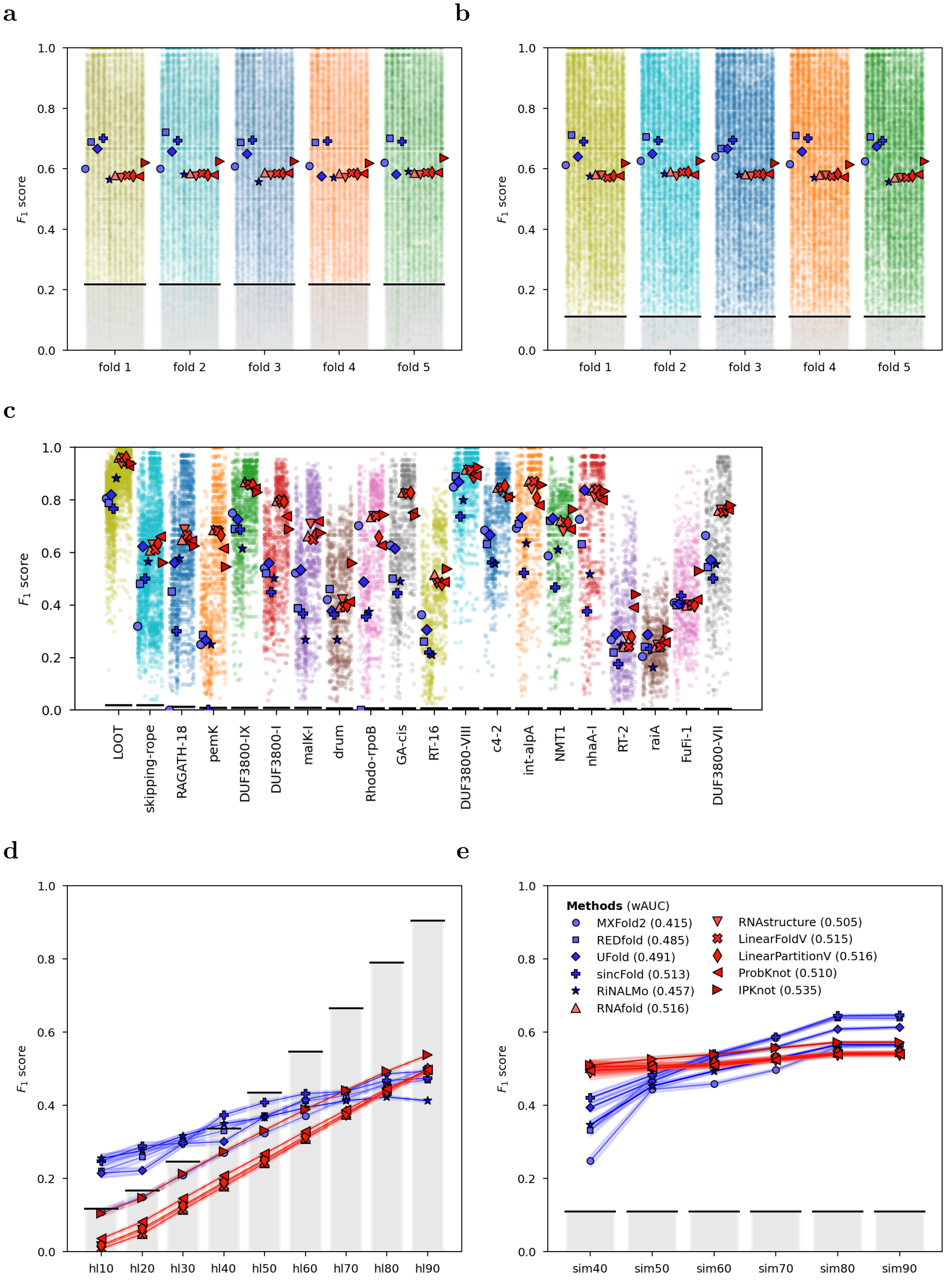


**Fig. S7.** Performance comparison for RNA secondary structure prediction methods according to different cross-validation strategies in the bpRNA dataset. Median $F_1$ for each prediction method is indicated with a different marker (DL prediction methods are depicted in blue, while classical prediction methods are reported in red). Gray bars show the proportion of test to train sequences in each fold. **a** $k$-fold, $F_1$ for each tested sequence is scattered in the background. **b** cluster-fold. **c** fam-fold. The test in this case has been done with the bpRNA-new subset. The 20 most numerous RNA families are shown. **d** hl-fold. **e** sim-fold. wAUC: area under the performance curve for each method, weighted by the difficulty of the partition.

### Supplementary tables

**Table S1.** Jensen-Shannon distance between distributions of test and train connection distances.

| RNA family | GU | AU | GC | All connections |
|---|---|---|---|---|
| **grp1** | 0.411 | 0.341 | 0.294 | 0.282 |
| **tmRNA** | 0.549 | 0.479 | 0.464 | **0.463** |
| **tRNA** | 0.644 | 0.644 | 0.639 | **0.632** |
| **5s** | 0.566 | 0.525 | 0.502 | **0.478** |
| **srp** | 0.516 | 0.448 | 0.406 | 0.404 |
| **telomerase** | 0.688 | 0.630 | 0.489 | 0.442 |
| **RNaseP** | 0.365 | 0.330 | 0.266 | 0.254 |
| **16s** | 0.513 | 0.399 | 0.323 | 0.296 |
| **23s** | 0.506 | 0.446 | 0.346 | 0.307 |

Jensen-Shannon distance between test and train distributions of connection distances for canonical pairs (GU, AU, GC and all of them) in reference structures, for each fold of the fam-fold in Archive II. Largest values in all connections are highlighted in bold.

**Table S2.** Detailed RNA family composition and performance at each hl-fold in Archive II.

| | | hl10 | hl20 | hl30 | hl40 | hl50 | hl60 | hl70 | hl80 | hl90 |
|---|---|---|---|---|---|---|---|---|---|---|
| | % test | 0.72 | 1.80 | 4.31 | 11.85 | 19.39 | 36.09 | 52.24 | 67.86 | 85.46 |
| **tRNA** | F1 RNAfold | 0.00 | 0.07 | 0.18 | 0.31 | 0.37 | 0.45 | 0.51 | 0.56 | 0.63 |
| | F1 sincFold | 0.65 | 0.63 | 0.77 | 0.87 | 0.87 | 0.89 | 0.91 | 0.89 | 0.86 |
| | % test | 0.78 | 2.81 | 9.20 | 16.99 | 27.44 | 38.11 | 51.52 | 72.49 | 92.21 |
| **5s** | F1 RNAfold | 0.02 | 0.13 | 0.21 | 0.25 | 0.29 | 0.34 | 0.42 | 0.52 | 0.59 |
| | F1 sincFold | 0.90 | 0.89 | 0.94 | 0.94 | 0.94 | 0.95 | 0.95 | 0.93 | 0.86 |
| | % test | | 3.68 | 22.29 | 45.45 | 79.22 | 97.62 | 99.78 | 99.78 | 99.78 |
| **tmRNA** | F1 RNAfold | | 0.12 | 0.24 | 0.31 | 0.38 | 0.41 | 0.42 | 0.42 | 0.42 |
| | F1 sincFold | | 0.57 | 0.71 | 0.63 | 0.58 | 0.36 | 0.28 | 0.29 | 0.32 |
| | % test | 0.88 | 3.30 | 7.71 | 16.08 | 34.80 | 68.50 | 91.41 | 99.34 | 100.00 |
| **RNaseP** | F1 RNAfold | 0.02 | 0.13 | 0.21 | 0.29 | 0.37 | 0.46 | 0.51 | 0.53 | 0.53 |
| | F1 sincFold | 0.26 | 0.60 | 0.59 | 0.61 | 0.70 | 0.73 | 0.62 | 0.39 | 0.39 |
| | % test | 6.43 | 11.87 | 19.17 | 25.16 | 30.83 | 43.14 | 58.17 | 74.29 | 88.56 |
| **srp** | F1 RNAfold | 0.01 | 0.06 | 0.12 | 0.18 | 0.23 | 0.31 | 0.40 | 0.49 | 0.55 |
| | F1 sincFold | 0.69 | 0.78 | 0.77 | 0.68 | 0.68 | 0.66 | 0.64 | 0.57 | 0.48 |
| | % test | 1.35 | 9.46 | 10.81 | 21.62 | 36.49 | 56.76 | 83.78 | 95.95 | 100.00 |
| **grp1** | F1 RNAfold | 0.00 | 0.14 | 0.17 | 0.26 | 0.35 | 0.42 | 0.49 | 0.53 | 0.54 |
| | F1 sincFold | 0.03 | 0.10 | 0.20 | 0.39 | 0.43 | 0.43 | 0.35 | 0.27 | 0.28 |
| | % test | | | | | 13.33 | 20.00 | 66.67 | 80.00 | 93.33 |
| **23s** | F1 RNAfold | | | | | 0.38 | 0.48 | 0.63 | 0.65 | 0.68 |
| | F1 sincFold | | | | | 0.56 | 0.51 | 0.41 | 0.31 | 0.35 |
| | % test | 4.55 | 13.64 | 19.70 | 27.27 | 37.88 | 57.58 | 65.15 | 80.30 | 95.45 |
| **16s** | F1 RNAfold | 0.00 | 0.12 | 0.18 | 0.23 | 0.29 | 0.37 | 0.38 | 0.45 | 0.51 |
| | F1 sincFold | 0.17 | 0.38 | 0.41 | 0.40 | 0.36 | 0.29 | 0.29 | 0.31 | 0.34 |
| | % test | | | 14.29 | 28.57 | 51.43 | 88.57 | 100.00 | 100.00 | 100.00 |
| **telomerase** | F1 RNAfold | | | 0.23 | 0.31 | 0.37 | 0.44 | 0.46 | 0.46 | 0.46 |
| | F1 sincFold | | | 0.68 | 0.67 | 0.61 | 0.44 | 0.11 | 0.13 | 0.18 |

For each RNA family in rows, the % of the family in the test partition, the $F_1$ score of a classical method (RNAfold), and the $F_1$ score of a DL method (sincFold) are reported for each fold of hl-fold in columns.

**Table S3.** Detailed analysis of the hl-folds in Archive II.

| | $N_{trn}$ | $N_{tst}$ | $\tilde{L}_{tr}$ | $\tilde{L}_{ts}$ | $\lvert\tilde{L}_{\Delta}\rvert$ | $GC^c_{tr}$ | $GC^c_{ts}$ | $JS_{GU}$ | $JS_{AU}$ | $JS_{GC}$ | $JS_{all}$ | $F_1^{DL}$ |
|---|---|---|---|---|---|---|---|---|---|---|---|---|
| hl10 | 3405 | 81 | 175 | 105 | 69.83 | 53.13 | 59.45 | 0.47 | 0.43 | 0.32 | 0.29 | 0.67 |
| hl20 | 3295 | 203 | 175 | 147 | 27.88 | 53.06 | 54.71 | 0.31 | 0.27 | 0.21 | 0.16 | 0.72 |
| hl30 | 3044 | 482 | 171 | 183 | 12.33 | 53.24 | 53.07 | 0.22 | 0.19 | 0.15 | 0.12 | 0.77 |
| hl40 | 2720 | 842 | 167 | 193 | 25.72 | 53.78 | 51.48 | 0.22 | 0.18 | 0.15 | 0.14 | 0.73 |
| hl50 | 2273 | 1339 | 157 | 203 | 45.16 | 54.44 | 50.88 | 0.26 | 0.21 | 0.18 | 0.18 | 0.73 |
| hl60 | 1712 | 1962 | 144 | 202 | 57.49 | 55.55 | 51.10 | 0.30 | 0.25 | 0.21 | 0.21 | 0.68 |
| hl70 | 1217 | 2512 | 130 | 196 | 66.03 | 56.55 | 51.51 | 0.31 | 0.28 | 0.24 | 0.23 | 0.66 |
| hl80 | 712 | 3073 | 120 | 187 | 66.95 | 56.91 | 52.33 | 0.32 | 0.30 | 0.26 | 0.25 | 0.63 |
| hl90 | 262 | 3573 | 104 | 179 | 74.32 | 58.57 | 52.82 | 0.44 | 0.37 | 0.34 | 0.32 | 0.61 |
| $\rho\ F_1$ | 0.77 | -0.77 | 0.82 | 0.11 | -0.94 | -0.82 | -0.11 | -0.74 | -0.76 | -0.87 | -0.93 | |

Rows: hl-folds and per-column correlation with $F_1$ score (at the last column). Columns: $N_{trn}$: number of training samples. $N_{tst}$: number of testing samples. $\tilde{L}_{tr}$: mean training length (lengths colored with respect to the maximum of 512 nts). $\tilde{L}_ts$: mean test length. $|\tilde{L}_{\Delta}|$: absolute difference between mean training and testing sequence lengths. GC content in training and testing sequences. Jensen-Shannon distance between test and train distributions of connection distances between canonical pairs (GU, AU, GC and all together) of reference structures. $F_1^{DL}$: performance of sincFold.
